# Prediction of exosomal miRNA-based biomarkers for liquid biopsy

**DOI:** 10.1101/2024.06.20.599824

**Authors:** Akanksha Arora, Gajendra Pal Singh Raghava

**Affiliations:** Department of Computational Biology, Indraprastha Institute of Information Technology, Okhla Phase 3, New Delhi-110020, India

**Keywords:** Exosomal miRNA, Exosome, Alignment-based approach, Artificial Intelligence based models, Liquid Biopsy, Biomarkers

## Abstract

In this study, we investigated the properties of exosomal miRNAs to identify potential biomarkers for liquid biopsy. We collected 956 exosomal and 956 non-exosomal miRNA sequences from RNALocate and miRBase to develop predictive models. Our initial analysis reveals that specific nucleotides are preferred at certain positions in miRNAs associated with exosomes. We employed an alignment-based approach, artificial intelligence (AI) models, and ensemble methods for predicting exosomal miRNAs. For the alignment-based approach, we used a motif-based method with MERCI and a similarity-based method with BLAST, achieving high precision but low coverage of about 29%. The AI models, developed using machine learning, deep learning techniques, and large language models, achieved a maximum AUC of 0.707 and an MCC of 0.268 on an independent dataset. Finally, our ensemble method, combining alignment-based and AI-based models, reached a maximum AUC of 0.73 and an MCC of 0.352 on an independent dataset. We have developed a web server, EmiRPred, to assist the scientific community in predicting and designing exosomal miRNAs and identifying associated motifs (https://webs.iiitd.edu.in/raghava/emirpred/).

**Key points:**

- Exosomal miRNAs have potential applications in liquid biopsy
- An ensemble method has been developed to predict and design exosomal miRNA
- An array of predictive models were built using alignment-based approaches and AI-based approaches (ML, DL, LLM)
- A variety of important features and motifs for exosomal miRNA have been identified
- A webserver, a python package, a github, and a standalone software have been created

## Introduction

Liquid biopsy represents a groundbreaking advancement in diagnostics, enabling the detection and monitoring of various diseases through the analysis of biofluids [1–3]. This minimally invasive technique permits sampling throughout the disease course, circumventing the risks associated with tissue biopsies [4]. Commonly utilized molecules in liquid biopsies include circulating tumor DNA (ctDNA), cell-free RNA (cfRNA), and circulating tumor cells (CTCs). However, these molecules face limitations, such as the lack of surface markers and the low abundance of the desired cell-free nucleic acids [5]. Recently, exosomes have emerged as promising candidates for liquid biopsy applications due to their stability and the critical roles they play in biological processes. Exosomes, small extracellular vesicles released by cells, carry a cargo of proteins, lipids, DNA, mRNA, miRNA, and metabolites [6,7]. Exosomal biomarkers are more stable than cell-free macromolecules and can reflect dynamic changes in the tumor microenvironment [8]. They can be detected earlier in the disease process, making them a more promising tool for early diagnosis and monitoring.

Exosomal miRNAs, which are small non-coding RNA molecules approximately 19–22 nucleotides long, play critical roles in cellular communication and the regulation of gene expression [9,10]. The biogenetic pathway of exosomal miRNAs initiates in the nucleus, where DNA sequences are transcribed by RNA polymerase to form primary miRNAs (pri-miRNAs). These pri-miRNAs adopt hairpin structures of 70–100 nucleotides following initial processing [11]. Exportin 5 mediates the transport of hairpin pri-miRNAs to the cytoplasm, where Dicer facilitates further processing [12,13]. Upon maturation, these double-stranded miRNAs are converted into single-stranded miRNAs and subsequently sorted into exosomes (see Figure 1). The incorporation of miRNAs into exosomes is a regulated process and is not random [14,15]. In the past, a number of miRNA-based biomarkers have been reported, such as miR-21, miR-1246, and miR-155 for non-small cell lung cancer, miR-17-5p, and miR-92a-3p for colorectal cancer, and miR-200b and miR-200c for ovarian cancer [16–18]. Beyond cancer, exosomal miRNAs can be used as potential biomarkers for other diseases, such as cardiovascular and neurological disorders, and personalized medicine.

In this study, first time a systematic attempt has been made to identify exosomal miRNAs aiming to fully leverage their potential in diagnostics and therapeutics. Firstly, we created a dataset of experimentally validated exosomal and non-exosomal miRNA sequences. Then, we divided this dataset into training and independent datasets, where the training dataset contains 80% miRNA sequences, and the validation/independent dataset contains the remaining 20% miRNA sequences. We trained and developed our models on the training dataset using five-fold cross-validation. All hyperparameters are tuned on the training dataset, and only the final model is evaluated on the validation dataset. This is important to avoid overoptimization of the models. We have tried all possible techniques to develop a method to predict exosomal miRNA with high precision. Initially, we used alignment-based approaches to predict exosomal miRNA. These approaches are based on motif search and sequence similarity search. These alignment-based approaches are only successful if the query sequence has a known motif or similarity with the annotated sequence; they fail in the absence of a motif or similarity. In order to overcome these limitations, we developed artificial intelligence (AI) models. Our AI-based models include machine learning, deep learning, and large language models. Finally, we developed an ensemble method that combines the strength of alignment and AI-based models. This comprehensive approach aims to predict exosomal miRNAs accurately, paving the way for their broader application in biomedical research and clinical practice.

**Figure 1:**
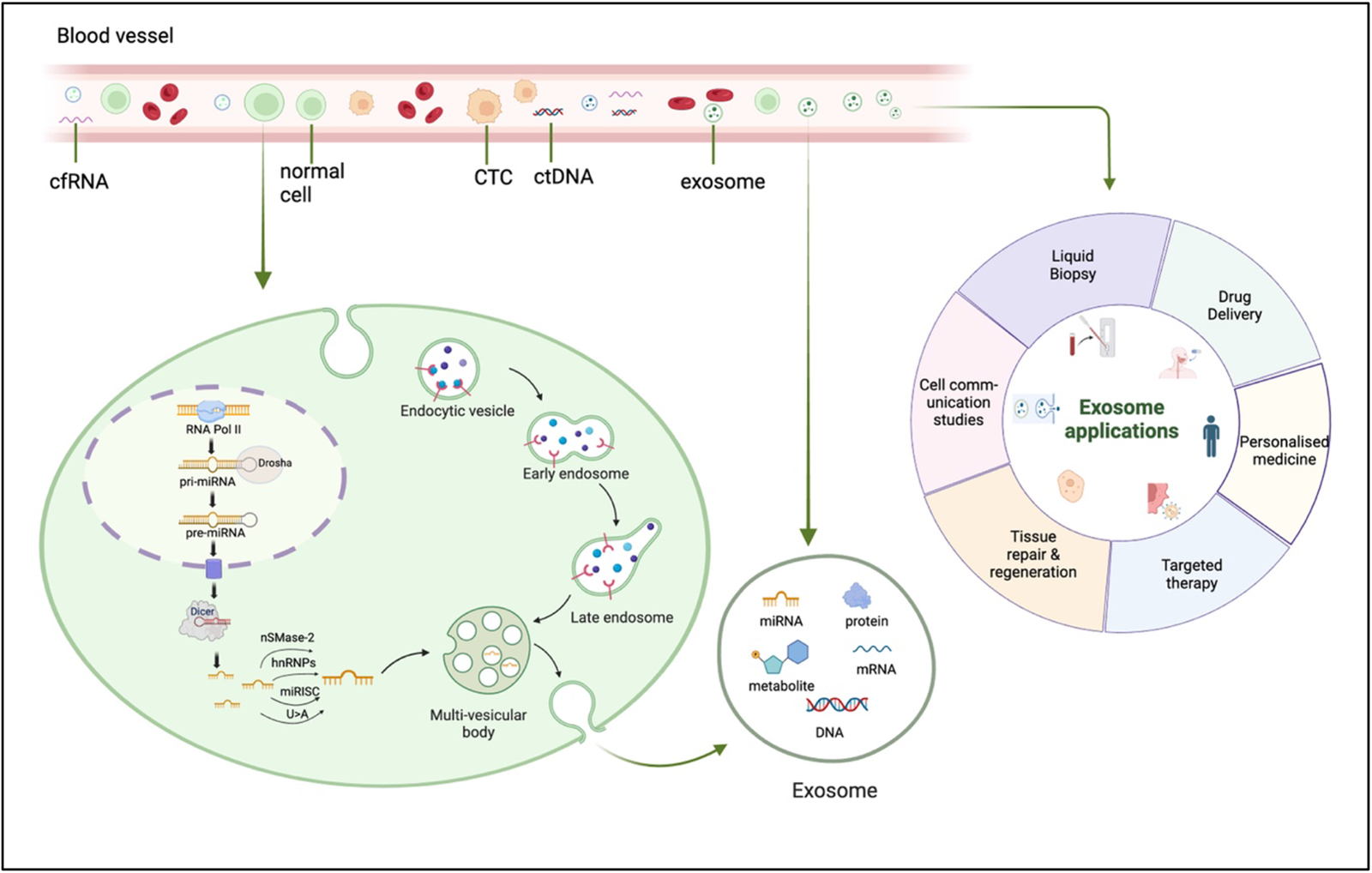
Shows biomolecules in body fluid commonly used in liquid biopsy, as well as the mechanism of secretion of miRNA from cell to exosomes.

## Methods

### 1. Data Collection and Preprocessing

In this study, datasets were creating using the data extracted from the databases: RNALocate and miRbase [19,20]. We collected the experimentally validated exosomal miRNA sequences from RNALocate. It contained about 1195 unique exosomal miRNA sequences for humans, with 956 mature miRNA sequences. Similarly, we collected experimentally validated miRNA sequences found in humans from RNALocate that were not detected in exosomes. Additionally, we sourced human miRNAs from miRBase and excluded those present in exosomes. This resulted in 1,694 unique non-exosomal miRNA sequences. We randomly selected 956 mature miRNAs from the non-exosomal miRNA to balance the dataset. Our final dataset contains 956 exosomal and 956 non-exosomal miRNAs. The sequence lengths ranged from 16 to 26 nucleotides, with most sequences being between 21 and 24 nucleotides long.

### 2. Alignment-based approaches

#### 2.1 Motif-Search

We used the MERCI (Motif Emerging with Classes Identification) tool to detect motifs found in exosomal miRNA sequences [21]. We identified motifs using the training dataset and then searched those motifs in the independent validation set. MERCI software has the option to select motifs that are exclusively present in the positive class (exosomal) and not present in the negative class (non-exosomal) from the training dataset. Additionally, this software offers a variety of options, such as the frequency of motifs and motifs with or without gaps.

#### 2.2 Similarity Search

In this study, we used blastn-short to annotate miRNA sequences based on their similarity to exosomal or non-exosomal miRNA sequences [22]. First, we built a database using the “makeblastdb” command for miRNA sequences present in the training dataset. To compute results for the training dataset, we removed self-hits and considered the top hit after removing self-hits. For the independent dataset, we considered the first hit to calculate results at various e-values. This method has been used in the previous studies [23,24].

### 3. AI-based classification methods

#### 3.1 Feature Generation

To develop a prediction model to predict exosomal and non-exosomal miRNA, we applied an array of techniques to extract meaningful features from miRNA sequences. They are discussed here:

##### 3.1.1 Composition-based Features

We utilized Nfeature to compute a wide range of composition-based features in miRNA sequences, including nucleotide composition, reverse complementary compositions, and auto-correlation [25]. Additionally, we computed features using Term Frequency-Inverse Document Frequency (TF-IDF), a statistical measure that evaluates the importance of a term in a document relative to a collection of documents. In our study, “terms” refer to k-mers, which are sequences of length k nucleotides, and a “document” refers to a sequence [26]. Term Frequency (TF) is the number of times a k-mer appears in a sequence, normalized by the total number of k-mers in the sequence. Inverse Document Frequency (IDF) is the logarithmically scaled inverse fraction of the sequences that contain the k-mer across the entire dataset. It measures how unique or rare a k-mer is within the whole dataset. By combining TF and IDF, TF-IDF assigns higher weights to k-mers that are frequent in a specific sequence but rare across the dataset, indicating that these k-mers are more informative and characteristic of the sequence’s content. The formulas for TF, IDF, and TF-IDF are explained in Equations 1, 2, and 3.

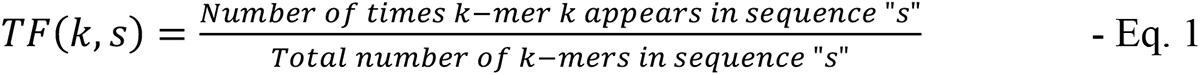

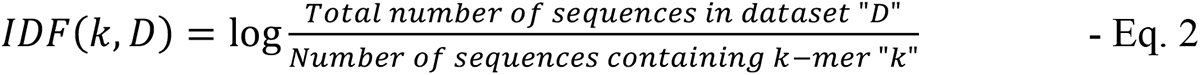

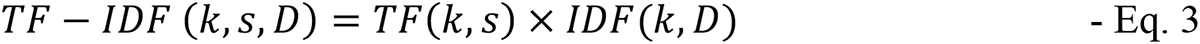

In Eq. 1,2, and 3, “k” represents a k-mer, “s” represents a sequence, and “D” represents the dataset of sequences.

The TFIDF features were computed for both the given sequences and reverse complementary sequences for kmer ranges from (1,1), (1,2),…(1,7). The best-performing TFIDF features were then also computed using different types of weighting – Term Frequency Collection (TFC) Weighting, Logarithmic Term Count (LTC) Weighting, and Entropy Weighting.

##### 3.1.2 Binary features

We computed binary profiles, or one-hot encoding features, for sequences to represent nucleotide sequences as binary vectors. To handle variable-length sequences, we use a technique called padding to standardize the length of the sequences. Since the maximum length of miRNA sequences is 26 nucleotides, each sequence is padded to a length of 26 by adding the dummy nucleotide ‘X’. For example, a sequence “AUTGTCGGGCUCUCCUAAUCU” of length 21 becomes “AUTGTCGGGCUCUCCUAAUCUXXXXX” of length 26. After padding, one-hot encoding features are computed by converting these sequences into binary profiles.

##### 3.1.3 Structure-based features

The structure-based features of miRNA sequences were computed using the RNAfold tool of Vienna RNA package 2.0, which predicts and identifies the RNA secondary structure based on the lowest possible free energy. It provides feature values for minimum free energy, ensemble free energy, centroid free energy, centroid diversity, the frequency of MFE structure in the ensemble, and ensemble diversity for each sequence in the dataset. In RNA folding, an ensemble typically refers to a collection of possible secondary structures that the RNA molecule can adopt [27]. The structures were predicted for each miRNA sequence in both exosomal and non-exosomal categories. These predicted structures were saved in .jpg image formats to develop further deep-learning classification models based on these structure images.

##### 3.1.4 Embeddings from Large Language Models

In this study, we generated embeddings using two types of BERT large language models (LLMs). BERT, developed by Google, is a transformer-based deep learning model. The following pre-trained models were used in this study.

a. BERT-Base Uncased: Uncased means that the text used during pre-training and fine-tuning is converted to lowercase, and no distinction is made between uppercase and lowercase letters. For example, “Sequence” and “sequence” would be treated as the same word. We extracted embeddings from this model after fine-tuning it on our training dataset [28].
b. DNABERT: DNABERT is a variant of BERT designed explicitly for processing DNA sequences. DNA-BERT is pre-trained on a large corpus of DNA sequences using self-supervised learning techniques. To use this model, we pre-processed our data and replaced “U” with “T” to suit the DNABERT model. We extracted the embeddings from this model by first fine-tuning the model on our sequence dataset [29].

#### 3.2 Prediction Models

##### 3.2.1 ML Models

We have used several ML algorithms to differentiate between exosomal and non-exosomal miRNA sequences. These algorithms involve K-Nearest Neighbours (KNN), Gaussian Naïve Bayes (GNB), Decision Tree (DT), Logistic Regression (LR), Extreme Gradient Boosting (XGB), Support Vector classifier (SVC), Extra Tree Classifier (ET) and Random Forest (RF) [30–37]. We have built prediction models using these ML algorithms on different sets of features defined in section 3.

##### 3.2.2 DL Models

We applied a deep learning (DL) classifier Convolutional Neural Networks (CNN) to classify exosomal and non-exosomal miRNA sequences. For the sequence classification task, we converted sequences to binary profiles as described in section 2.1. These binary profile of miRNA is classified using CNN algorithm. Additionally, two deep learning algorithms, CNN and ResNet (Residual Network), were applied to the miRNA structure images generated as described in section 2.3.2. ResNet utilizes shortcut connections to facilitate residual learning, enabling deep networks to train effectively. We used the ResNet50 variant, a 50-layer version of the ResNet architecture, as it balances complexity and efficiency [38–42].

##### 3.2.3 LLM Models

Large Language Models (LLMs) excel in text classification due to their deep contextual understanding. They are pre-trained on extensive corpora and can be fine-tuned for specific tasks like sequence classification in our study [43]. They adapt well to various domains. The pre-training in LLMs provides efficient feature extraction, enabling accurate classifications even with smaller datasets. We have used two types of LLM models in our study to classify exosomal and non-exosomal miRNA sequences. We have used BERT-based uncased and DNABERT, as mentioned in section 2.5, and fine-tuned them according to our sequence length and dataset composition. These fine-tuned models were then used to make predictions on our dataset and classify the sequences as exosomal and non-exosomal.

#### 3.3 Cross-validation and performance metrics

The entire dataset, consisting of 1912 sequences, was divided into an 80:20 ratio, with 80% used for training and 20% for validation. A five-fold cross-validation technique was employed on 80% of the training data to evaluate the machine learning (ML) and deep learning (DL) models, keeping the remaining 20% unseen for the models. In five-fold cross-validation, the training data is split into five parts: four parts for training and one part for internal validation. This process is repeated five times, ensuring each fold serves as the test set once. The models were assessed using both threshold-dependent and threshold-independent metrics. Evaluation metrics included sensitivity, specificity, Matthews correlation coefficient (MCC), accuracy, and Area Under the Receiver Operating Characteristics (AUROC). AUROC is threshold-independent, while the other metrics depend on the threshold. Specificity, sensitivity, and MCC were optimized for the best threshold values. These metrics have been used in previous studies to measure the performance of models [44–47]

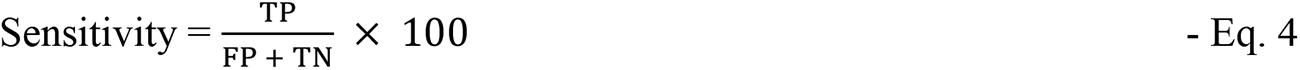

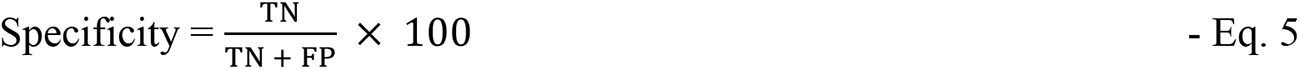

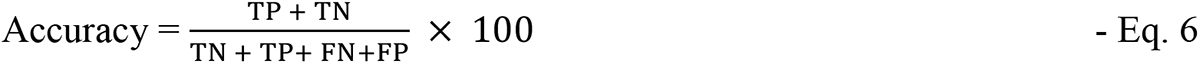

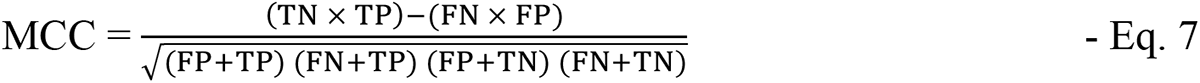

Where TP, FP, TN, and FN are true positive, false positive, true negative, and false negative, respectively.

### 4. Ensemble method

We attempted to improve the prediction of our best-performing prediction model developed on the best set of features by using a combination of various findings. This approach applies a weighted scoring method that combines three techniques: (i) motif-based approach, (ii) similarity search using BLAST, and (iii) ML/DL prediction based methods. In this approach, we assign a score of +0.5 to an miRNA sequence if it has an exosomal motif and 0 if no exosomal motif is found. We further add +0.5 if the sequence is found to be similar to an exosomal miRNA sequence using the BLAST algorithm. These scores are then combined with best performing ML/DL model prediction scores, obtained via the predict_proba() function in scikit-learn, which gives the probability of a sequence belonging to a specific class instead of a binary outcome [48]. Together, the motif score, similarity search score, and best ML/DL model score provide a combined score for each sequence, ranging from 0 to 2 as shown in Eq. 8 and 9. By analyzing these overall scores, the sequences were classified as either exosomal or non-exosomal. Several studies have previously utilized this hybrid/ensemble approach [7,49].

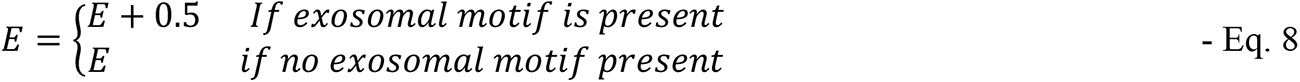

Here, E = Prediction probability score obtained from best performing ML/DL model and *E’*= Score obtained after adding scores from the motif-based approach

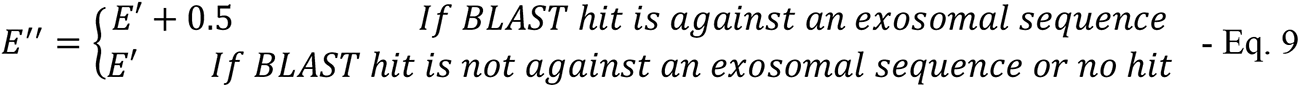

Here, *E”* = Final score obtained from the best performing ML/DL model, motif-based approach, and BLAST-based approach ranging from 0 to 2

## Results

In this study, we employed different techniques to predict exosomal miRNA, which can be categorized into three main categories: i) Alignment-based approaches, ii) AI-based models, and iii) Ensemble methods. Alignment-based approaches encompass motif-search using MERCI software and similarity-search using BLAST. AI-based methods involve the development of machine learning (ML), deep learning (DL), and large language model (LLM) based models. The AI-based models used numerous features and their combinations, such as compositional, binary, structural features, and embeddings. To harness the full potential of alignment-based and AI-based techniques, we devised an ensemble method that integrates the strengths of both approaches. The comprehensive architecture of EmiRPred and the methodologies employed are illustrated in Figure 2.

**Figure 2:**
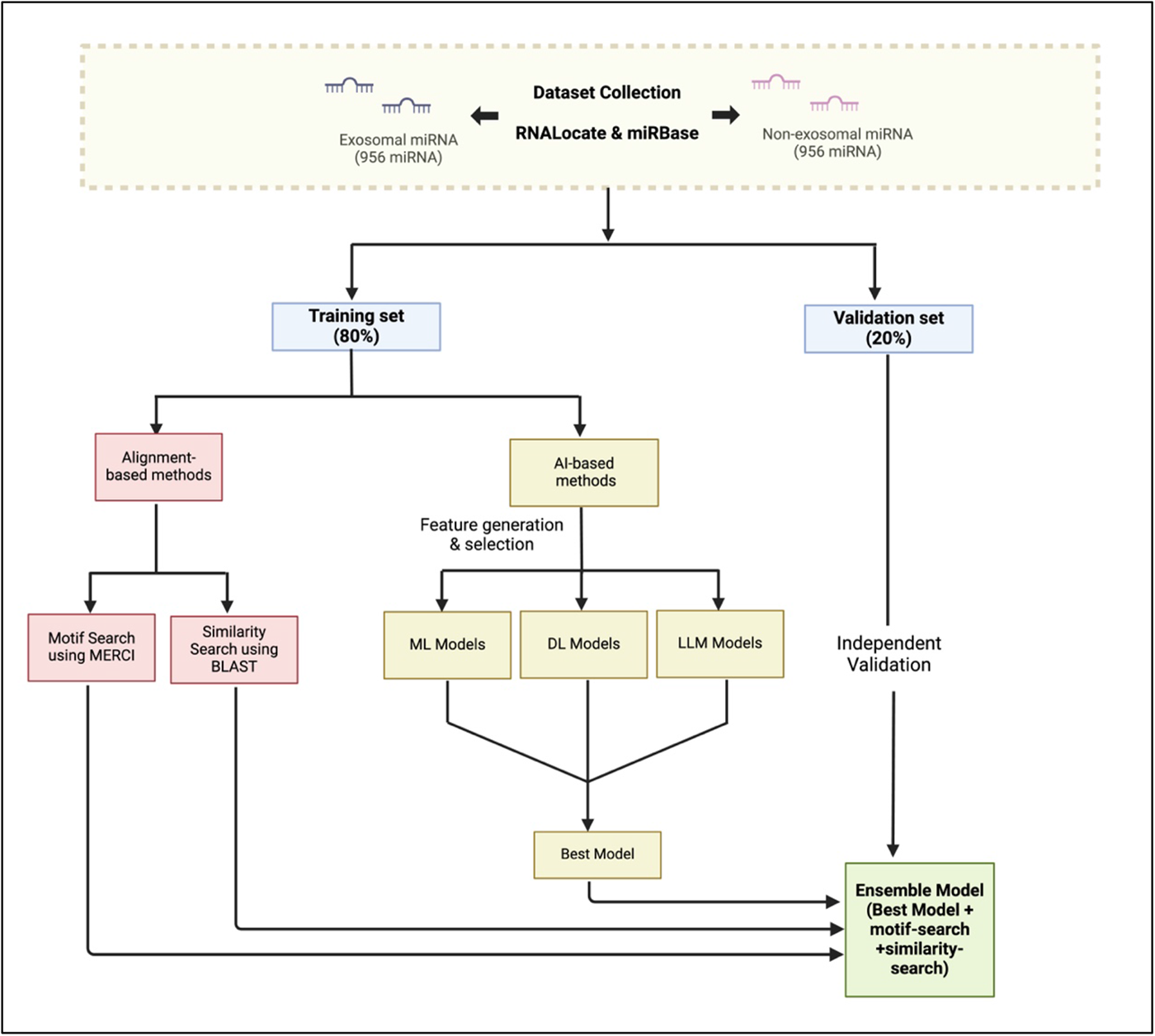
The complete architecture of algorithm used in EmiRPred

### 1. Alignment-based classification methods

#### 1.1 Motif-Search

In our study, we discovered motifs within exosomal miRNA using MERCI software applied with various parameters [21]. For instance, employing a Gap value of 0 led to the discovery of 11 motifs that covered 26 exosomal sequences in the training dataset and 8 sequences in the validation dataset. Similarly, applying a Gap value of 1 uncovered 11 motifs spanning 30 exosomal sequences in the training dataset and 10 sequences in the validation dataset. The detailed results of motif discovery using MERCI under different settings are presented in Table 1, along with the sequences covered in the validation dataset.

**Table 1:**
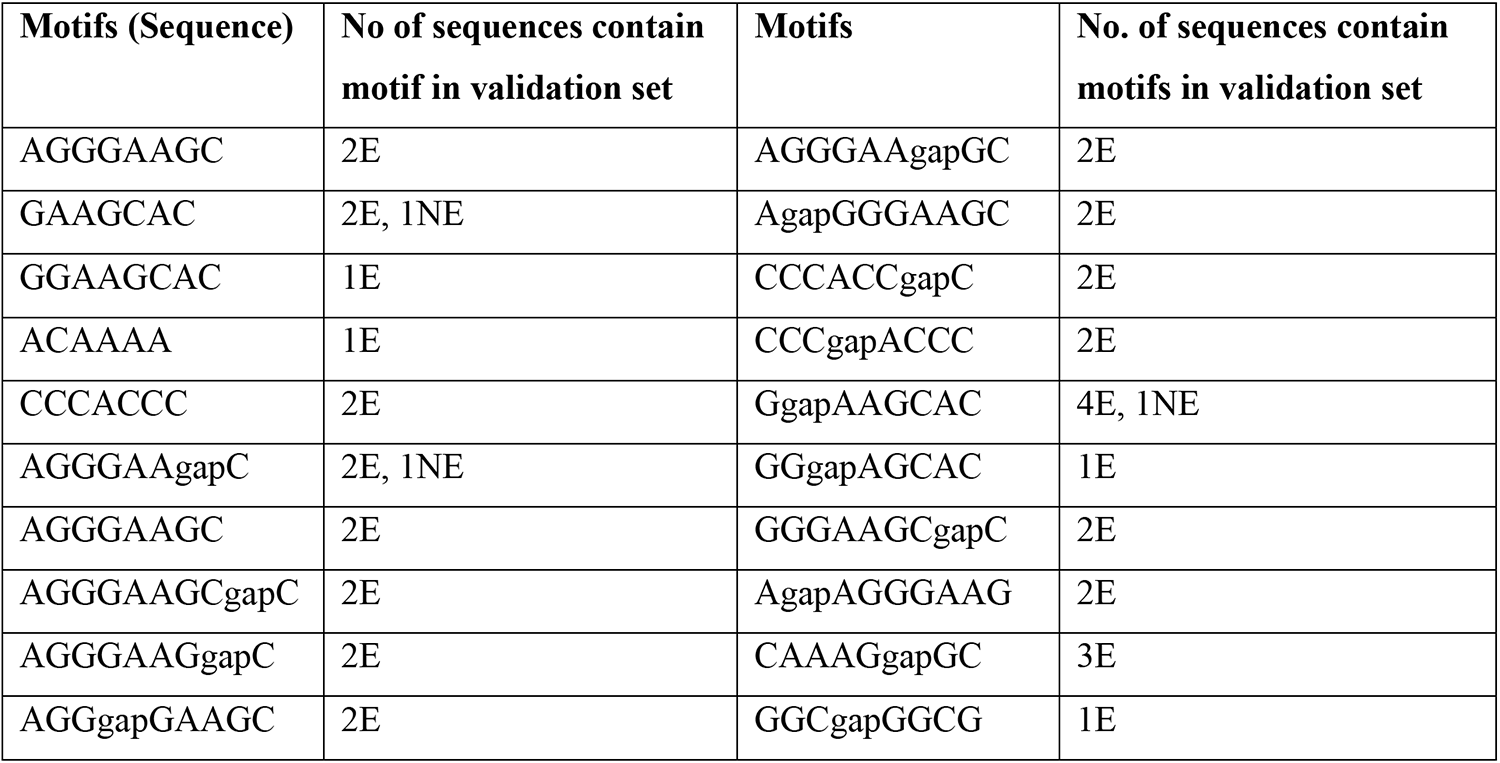
List of exosomal motifs discovered in the training set and their occurrence in independent/validation dataset (E for exosomal, NE for non-exosomal)

#### 1.2 Similarity-search using BLAST

In this paper, we applied blastn-short to perform a similarity search against a training dataset of exosomal and non-exosomal miRNA for the e-values ranging from 10^-6^ to 10^6^. [22]. Using this approach, we obtained an optimal performance at e-value 10^-2^, with 233 correct hits and 85 incorrect hits for exosomal miRNA sequences in the training set; and 66 correct hits and 24 incorrect hits for exosomal miRNA sequences in the validation set. The e-values lesser than 10^-2^ did not show enough coverage for sequences, and the values greater than 10^-2^ showed a higher error rate. The full results for BLAST from e-values 10^-6^ to 10^6^ are shown in Table 2.

**Table 2:** Number of correct, incorrect, and total hits for exosomal miRNA sequences in training and validation set for e-values ranging from 10^-6^ to 10^6^.

| e-value | Training Dataset |  |  |  | Validation set |  |  |  |
| --- | --- | --- | --- | --- | --- | --- | --- | --- |
|  | Correct hits | Incorrect hits | No hits | Total | Correct hits | Incorrect hits | No hits | Total |
| $10^6$ | 769 | 758 | 0 | 1527 | 185 | 198 | 0 | 383 |
| $10^5$ | 769 | 758 | 0 | 1527 | 185 | 198 | 0 | 383 |
| $10^4$ | 769 | 758 | 0 | 1527 | 185 | 198 | 0 | 383 |
| $10^3$ | 769 | 758 | 0 | 1527 | 185 | 198 | 0 | 383 |
| $10^2$ | 769 | 758 | 0 | 1527 | 185 | 198 | 0 | 383 |
| $10^1$ | 769 | 758 | 0 | 1527 | 185 | 198 | 0 | 383 |
| $10^0(1)$ | 626 | 559 | 342 | 1527 | 152 | 146 | 85 | 383 |
| $10^{-1}$ | 300 | 143 | 1084 | 1527 | 79 | 44 | 260 | 383 |
| $10^{-2}$ | 233 | 85 | 1209 | 1527 | 66 | 24 | 293 | 383 |
| $10^{-3}$ | 175 | 53 | 1299 | 1527 | 49 | 15 | 319 | 383 |
| $10^{-4}$ | 119 | 34 | 1374 | 1527 | 35 | 13 | 335 | 383 |
| $10^{-5}$ | 99 | 24 | 1404 | 1527 | 30 | 11 | 342 | 383 |
| $10^{-6}$ | 50 | 14 | 1463 | 1527 | 14 | 6 | 363 | 383 |

### 2. AI-based classification methods

#### 2.1 ML Models

Initially, we computed a wide range of sequence features for miRNA sequences; details are given in the Materials & Methods section. Next, we applied a variety of machine learning techniques for developing prediction models that includes DT, KNN, XGB, LR, SVC, RF, and ET (Figure 3, Table S1). were applied. The performances of ML-based models developed using different classes of features are as follows:

##### 2.1.1 Composition-based Features

We calculated various composition-based features using Nfeature, including nucleotide composition, composition of reverse complement, nucleotide repeat index, distance distribution of nucleotides, and pseudo composition [25]. Our RF-based model developed using the composition of reverse complement RNA sequence for k-mers 3 and 4 (RDK-3 & RDK-4) achieved a maximum AUC of 0.677. Our KNN model developed using TF-IDF achieved a maximum AUC of 0.656.

##### 2.1.2 Binary profile

We computed binary profiles or one hot encoding features to represent the nucleotide sequences as binary vectors. We developed machine learning-based models using these binary profiles. An SVM-based model performed the best on these features with an AUC of 0.642, on the validation dataset.

##### 2.1.3 Secondary Structure-based features

We estimated secondary structure-based features for miRNA sequences using the RNAfold tool from the ViennaRNA package 2.0 [27]. These features included minimum free energy, ensemble free energy, centroid free energy, centroid diversity, frequency of the minimum free energy (MFE) structure in the ensemble, and ensemble diversity. A logistic regression model, trained on these secondary structure-based features, achieved the highest AUC of 0.558 on the validation dataset.

##### 2.1.4 Embeddings from Large Language Model

To extract the embeddings from LLM models, we used two pre-trained models – DNABert and BERT-base-uncased and then fine-tuned them on our training dataset [28,29]. The embeddings were then extracted from the fine-tuned models for miRNA sequences which were then used to develop models using machine learning models. A random forest model achieved the maximum AUC 0.598 and 0.565 using embeddings of DNABERT and BERT-base-uncased, respectively.

##### 2.1.5 Best Features

We developed machine learning models using a combination of best features, including mononucleotide binary profiles, the composition of reverse complementary miRNA sequences (RDK-3, RDK-4), and TF-IDF. This resulted in a total of 382 features, which were normalized using the StandardScaler from the Scikit-learn package [48]. Our Extra Trees model, built using these features, achieved an AUC of 0.707 on the validation dataset. Additionally, a Mann-Whitney test was conducted on the 382 features to identify those that significantly differentiate exosomal from non-exosomal miRNA sequences. We found 75 features with p-values less than 0.05, indicating a significant difference between the two classes. The results for the individual best features and the combined best features are provided in Table 3 and Table 4, respectively. The results for the Mann-Whitney test performed on these features in given in Supplementary Table S2.

##### 2.1.6 Feature Importance

We selected the top 20 features from the best-performing set of features according to their feature importance in the model. It was observed that 17 out of the 20 features were significantly different (p-value <0.05) as calculated by the Mann-Whitney test in Section 3.6 of “Results”. In binary profile features, it was observed that C at the 1^st^ position (C_1), U at the 21^st^ position (U_21), and G at the 15^th^ position (G_15) are seen in more exosomal miRNA sequences than non-exosomal miRNA sequences. From reverse complement sequence compositional features (RDKs), it is seen that RDK_CAC is significantly increased in exosomal miRNA sequences than in non-exosomal sequences (p-value<0.0001). In TFIDF features from the range of (1,3) in reverse complement miRNA sequences, it is seen that “U” was found in abundance in exosomal sequences than non-exosomal, dinucleotides “AC” and “UC”, and trinucleotides “GGA”, “GGC”, and “GUC” were also significantly increased in exosomal sequences. The top 20 most important features in distinguishing exosomal and non-exosomal miRNA sequences have been shown in Figure 5. The results for the importance of all features are given in Supplementary Table S3.

#### 2.2 DL Models

##### 2.2.1 Sequences

We applied the CNN algorithm to develop a prediction model using the binary profiles of miRNA sequences [38]. The model gave an AUC of 0.611 on the training dataset after five-fold cross-validation and 0.621 on an independent validation dataset. The detailed results for CNN model applied on miRNA sequences are given in Supplementary Table S1.

##### 2.2.2 Structure Images

We obtained the secondary structures of miRNA sequences using the RNAfold of the Vienna RNA package [27]. RNAfold predicted secondary structure of miRNA in form of images. These images were used to develop models CNN and ResNet 50 [40,42]. Our CNN and ResNet50 achieved AUCs of 0.551 and 0.553, respectively. The results for these models applied on RNA secondary structure images are given in Supplementary Table S1.

#### 2.3 LLM Models

We used pre-trained LLM models like DNABERT and BERT-base-uncased and fine-tuned them on our data [28,29]. DNABERT was able to differentiate between the exosomal and non-exosomal sequences with an AUC of 0.535 and BERT-base-uncased was able to differential them with an AUC of 0.608. The detailed results for finetuned LLM models are given in Supplementary Table S1.

**Table 3:** AI-based methods results for the best performing features.

| Binary features / One hot encoding - mononucleotides |  |  |  |  |  |  |  |  |  |  |  |
| --- | --- | --- | --- | --- | --- | --- | --- | --- | --- | --- | --- |
|  |  | Training set |  |  |  |  | Validation set |  |  |  |  |
| Model | Thr | Sens | Spec | Acc | AUC | MCC | Sens | Spec | Acc | AUC | MCC |
| DT | 0.51 | 54.09 | 54.49 | 54.28 | 0.553 | 0.086 | 58.38 | 53.03 | 55.61 | 0.571 | 0.114 |
| RF | 0.50 | 61.61 | 59.63 | 60.63 | 0.640 | 0.212 | 58.92 | 63.64 | 61.36 | 0.634 | 0.226 |
| LR | 0.50 | 54.99 | 54.35 | 54.68 | 0.578 | 0.093 | 52.43 | 60.10 | 56.40 | 0.560 | 0.126 |
| XGB | 0.50 | 57.20 | 57.12 | 57.16 | 0.614 | 0.143 | 56.76 | 52.02 | 54.31 | 0.571 | 0.088 |
| KN | 0.51 | 55.51 | 59.76 | 57.62 | 0.613 | 0.153 | 58.92 | 60.10 | 59.53 | 0.614 | 0.190 |
| GNB | 0.84 | 49.94 | 49.47 | 49.71 | 0.489 | -0.006 | 35.14 | 74.24 | 55.35 | 0.555 | 0.102 |
| SVC | 0.50 | 59.40 | 60.95 | 60.17 | 0.634 | 0.204 | 61.08 | 62.12 | 61.62 | 0.642 | 0.232 |
| ET | 0.50 | 56.42 | 59.50 | 57.95 | 0.628 | 0.159 | 58.38 | 65.66 | 62.14 | 0.633 | 0.241 |
| Reverse Complement Sequence Composition (RDk3 + RDk 4) |  |  |  |  |  |  |  |  |  |  |  |
|  |  | Training set |  |  |  |  | Validation set |  |  |  |  |
| Model | Thr | Sens | Spec | Acc | AUC | MCC | Sens | Spec | Acc | AUC | MCC |
| DT | 0.53 | 54.86 | 49.21 | 52.06 | 0.529 | 0.041 | 58.92 | 52.02 | 55.35 | 0.559 | 0.110 |
| RF | 0.50 | 58.89 | 56.99 | 57.95 | 0.627 | 0.159 | 61.62 | 63.64 | 62.66 | 0.677 | 0.253 |
| LR | 0.50 | 57.07 | 56.60 | 56.84 | 0.596 | 0.137 | 56.22 | 60.10 | 58.23 | 0.602 | 0.163 |
| XGB | 0.50 | 58.50 | 56.73 | 57.62 | 0.601 | 0.152 | 58.38 | 58.59 | 58.49 | 0.620 | 0.170 |
| KN | 0.48 | 60.31 | 57.52 | 58.93 | 0.624 | 0.178 | 65.41 | 53.54 | 59.27 | 0.644 | 0.191 |
| GNB | 0.08 | 55.90 | 56.46 | 56.18 | 0.594 | 0.124 | 52.97 | 55.05 | 54.05 | 0.568 | 0.080 |
| SVC | 0.49 | 40.08 | 58.44 | 49.18 | 0.485 | -0.015 | 100.0<br>0 | 0.00 | 48.30 | 0.360 | 0.000 |
| ET | 0.50 | 59.66 | 59.37 | 59.52 | 0.639 | 0.190 | 62.70 | 59.09 | 60.84 | 0.662 | 0.218 |
| Term Frequency- Inverse Document Frequency (TFIDF)<br>on reverse complementary sequence (kmer range – 1,3) |  |  |  |  |  |  |  |  |  |  |  |
|  |  | Training set |  |  |  |  | Validation set |  |  |  |  |
| Model | Thr | Sensitivity | Specificity | Acc | AUC | MCC | Sensitivity | Specificity | Acc | AUC | MCC |
| DT | 0.01 | 55.25 | 55.28 | 55.26 | 0.553 | 0.105 | 52.43 | 60.10 | 56.40 | 0.563 | 0.126 |
| RF | 0.51 | 59.53 | 57.78 | 58.67 | 0.622 | 0.173 | 63.78 | 56.57 | 60.05 | 0.632 | 0.204 |
| LR | 0.51 | 54.86 | 56.86 | 55.85 | 0.594 | 0.117 | 51.89 | 53.03 | 52.48 | 0.565 | 0.049 |
| XGB | 0.51 | 58.75 | 58.58 | 58.67 | 0.605 | 0.173 | 62.70 | 53.03 | 57.70 | 0.620 | 0.158 |
| KN | 0.41 | 56.94 | 59.37 | 58.14 | 0.602 | 0.163 | 71.35 | 58.59 | 64.75 | 0.656 | 0.301 |
| GNB | 0.46 | 56.81 | 56.73 | 56.77 | 0.588 | 0.135 | 52.43 | 51.52 | 51.96 | 0.564 | 0.039 |
| SVC | 0.51 | 57.85 | 62.01 | 59.91 | 0.635 | 0.199 | 61.08 | 61.11 | 61.10 | 0.634 | 0.222 |
| ET | 0.51 | 61.61 | 62.01 | 61.81 | 0.236 | 0.652 | 58.92 | 58.59 | 58.75 | 0.629 | 0.175 |

**Table 4:** Results for a) best performing features combined (AI-based methods), and b) hybrid model: best-performing features (AI-based methods) combined with motif-search and similarity search (Alignment-based methods)

| Best features combined – |  |  |  |  |  |  |  |  |  |  |  |
| --- | --- | --- | --- | --- | --- | --- | --- | --- | --- | --- | --- |
| One hot encoding (mononucleotides) + RDK-3 + RDK-4 + Reverse complement TFIDF (1,3) |  |  |  |  |  |  |  |  |  |  |  |
|  |  | Training set |  |  |  |  | Validation set |  |  |  |  |
| Model | Thr | Sens | Spec | Acc | AUC | MCC | Sens | Spec | Acc | AUC | MCC |
| DT | 0.50 | 56.16 | 53.83 | 55.00 | 0.550 | 0.100 | 56.76 | 54.55 | 55.61 | 0.557 | 0.113 |
| RF | 0.51 | 57.59 | 61.74 | 59.65 | 0.648 | 0.193 | 60.54 | 59.09 | 59.79 | 0.637 | 0.196 |
| LR | 0.50 | 57.46 | 56.99 | 57.23 | 0.602 | 0.144 | 60.54 | 58.59 | 59.53 | 0.620 | 0.191 |
| XGB | 0.55 | 55.12 | 63.98 | 59.52 | 0.636 | 0.192 | 65.41 | 56.57 | 60.84 | 0.640 | 0.220 |
| KN | 0.50 | 57.98 | 59.63 | 58.80 | 0.621 | 0.176 | 63.24 | 58.08 | 60.57 | 0.656 | 0.213 |
| GNB | 0.98 | 55.25 | 53.83 | 54.55 | 0.558 | 0.091 | 38.38 | 70.71 | 55.09 | 0.586 | 0.096 |
| SVC | 0.51 | 55.77 | 62.80 | 59.25 | 0.644 | 0.186 | 64.32 | 64.65 | 64.49 | 0.678 | 0.290 |
| ET | 0.50 | 62.39 | 62.01 | 62.20 | 0.672 | 0.244 | 63.24 | 63.64 | 63.45 | 0.707 | 0.269 |
| Hybrid model: Best features combined with motif-search and similarity search |  |  |  |  |  |  |  |  |  |  |  |
|  |  | Training set |  |  |  |  | Validation set |  |  |  |  |
| Model | Thr | Sens | Spec | Acc | AUC | MCC | Sens | Spec | Acc | AUC | MCC |
| DT | 0.51 | 57.33 | 53.83 | 55.59 | 0.614 | 0.112 | 57.84 | 54.04 | 55.87 | 0.622 | 0.119 |
| RF | 0.55 | 61.22 | 65.83 | 63.51 | 0.694 | 0.271 | 64.86 | 62.63 | 63.71 | 0.685 | 0.275 |
| LR | 0.59 | 60.96 | 60.16 | 60.56 | 0.663 | 0.211 | 64.32 | 61.62 | 62.92 | 0.687 | 0.259 |
| XGB | 0.59 | 61.74 | 61.21 | 61.48 | 0.680 | 0.230 | 69.19 | 53.54 | 61.10 | 0.694 | 0.230 |
| KN | 0.54 | 69.00 | 54.35 | 61.74 | 0.673 | 0.236 | 70.81 | 54.04 | 62.14 | 0.690 | 0.252 |
| GNB | 0.93 | 60.44 | 50.00 | 55.26 | 0.618 | 0.105 | 49.73 | 67.68 | 59.01 | 0.646 | 0.177 |
| SVC | 0.54 | 60.44 | 62.27 | 61.35 | 0.691 | 0.227 | 67.03 | 65.15 | 66.06 | 0.711 | 0.322 |
| ET | 0.52 | 65.63 | 62.93 | 64.29 | 0.703 | 0.286 | 67.57 | 67.68 | 67.62 | 0.730 | 0.352 |

**Figure 3:**
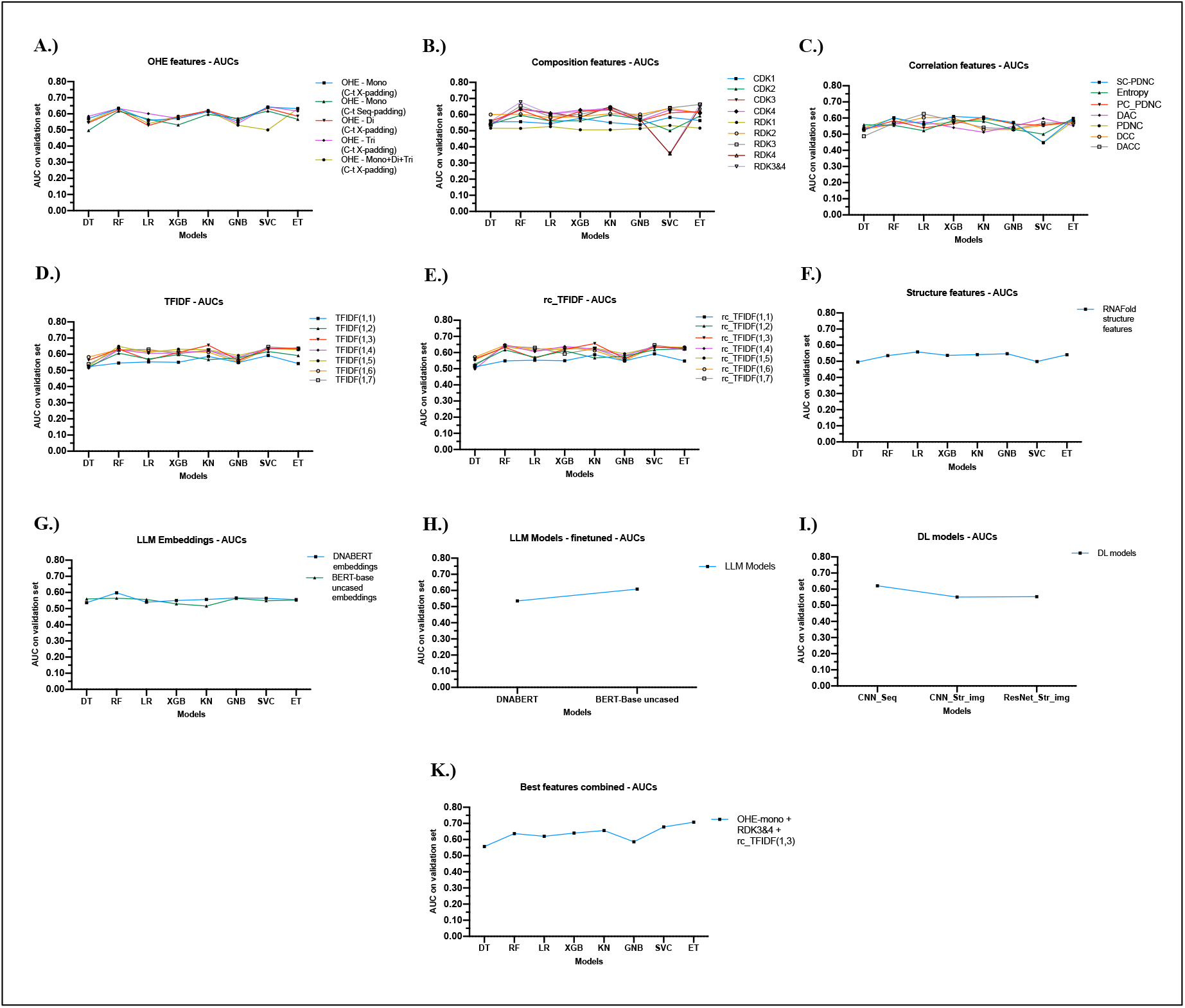
Comparisons of AUCs for different approaches: A) One hot encoding (OHE) features, B) Composition features, C) Correlation features, D) Term frequency-inverse document frequency (TFIDF) for different k-mer ranges, E) Term frequency – inverse document frequency for reverse complementary sequences (rc_TFIDF) for different k-mer ranges, F) Structural features extracted using RNAFold, G) LLM Embeddings of DNABERT, and BERT-base-uncased, H) Finetuned LLM models – DNABERT and BERT-base-uncased, I) DL models: CNN on sequence, CNN on structure images, ResNet50 on structure images J) Best features combined (RDK-3, RDK-4, rc_TFIDF (1,3), and OHE mononucleotides)

### 3. Hybrid classification method

To develop a model that is able to differentiate between exosomal and non-exosomal miRNA sequence classes with high accuracy, we developed a hybrid model that combined the predictive power of our best performing ML model on the best identified features with motif-search, and similarity search algorithm. The best performing prediction model is ET model with an AUC of 0.707 on independent validation dataset, which was improved to 0.712 when motif-search algorithm results were added using the hybrid approach scoring system described in “Methods”. The AUC further improved to 0.73 when we added the similarity-search algorithm using BLAST on e-value = 10^-2^ to our model. Along with the AUC of 0.73, this final model also showed an accuracy of 67.62% and an MCC of 0.352 on an independent validation set. The resulting metrics for both training and independent validation set for this hybrid model are given in Table 4. The Area Under the Receiver Operating Curves (AUROC) for the training and validation set for the hybrid model are shown in Figure 4.

**Figure 4:**
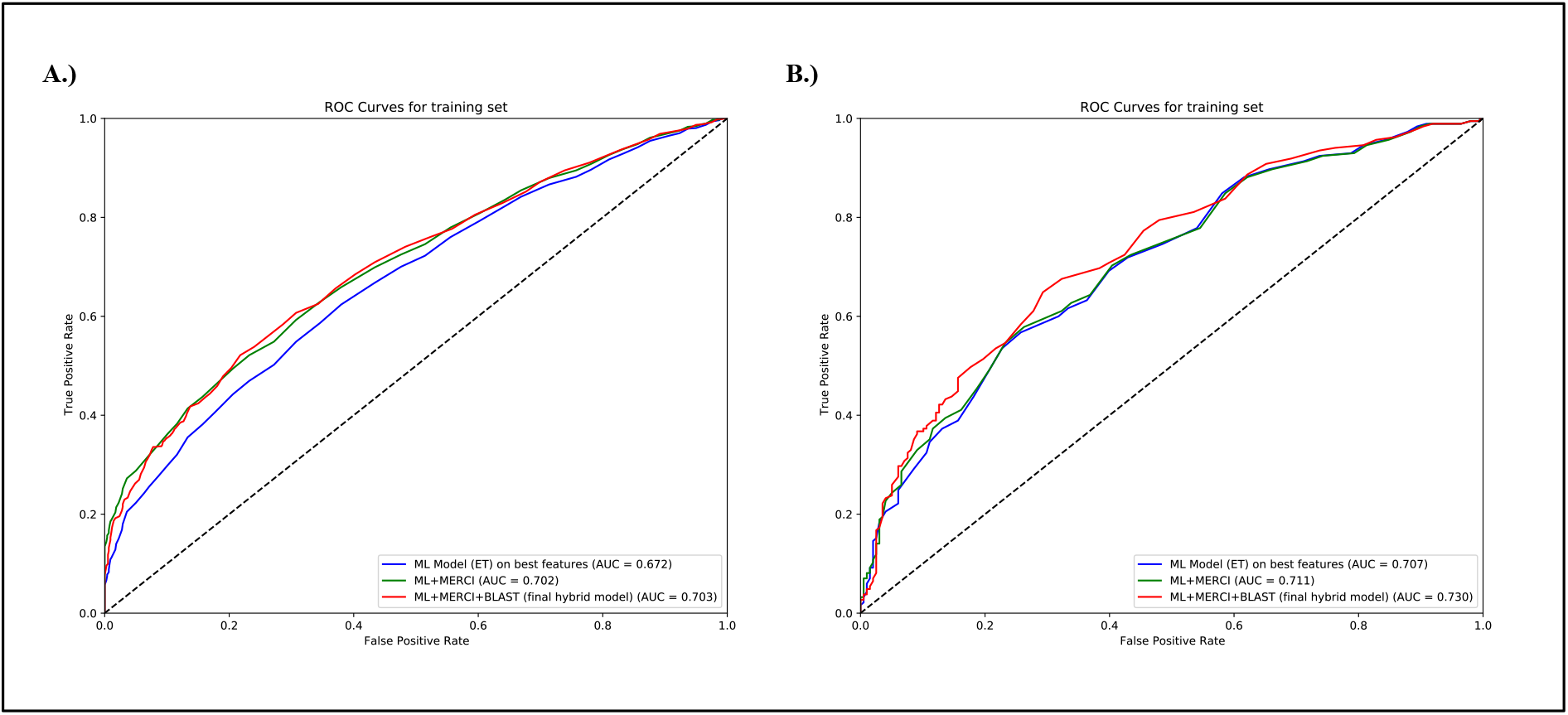
AUROC graphs for the hybrid model A) Training set and B) Independent Validation set

**Figure 5:**
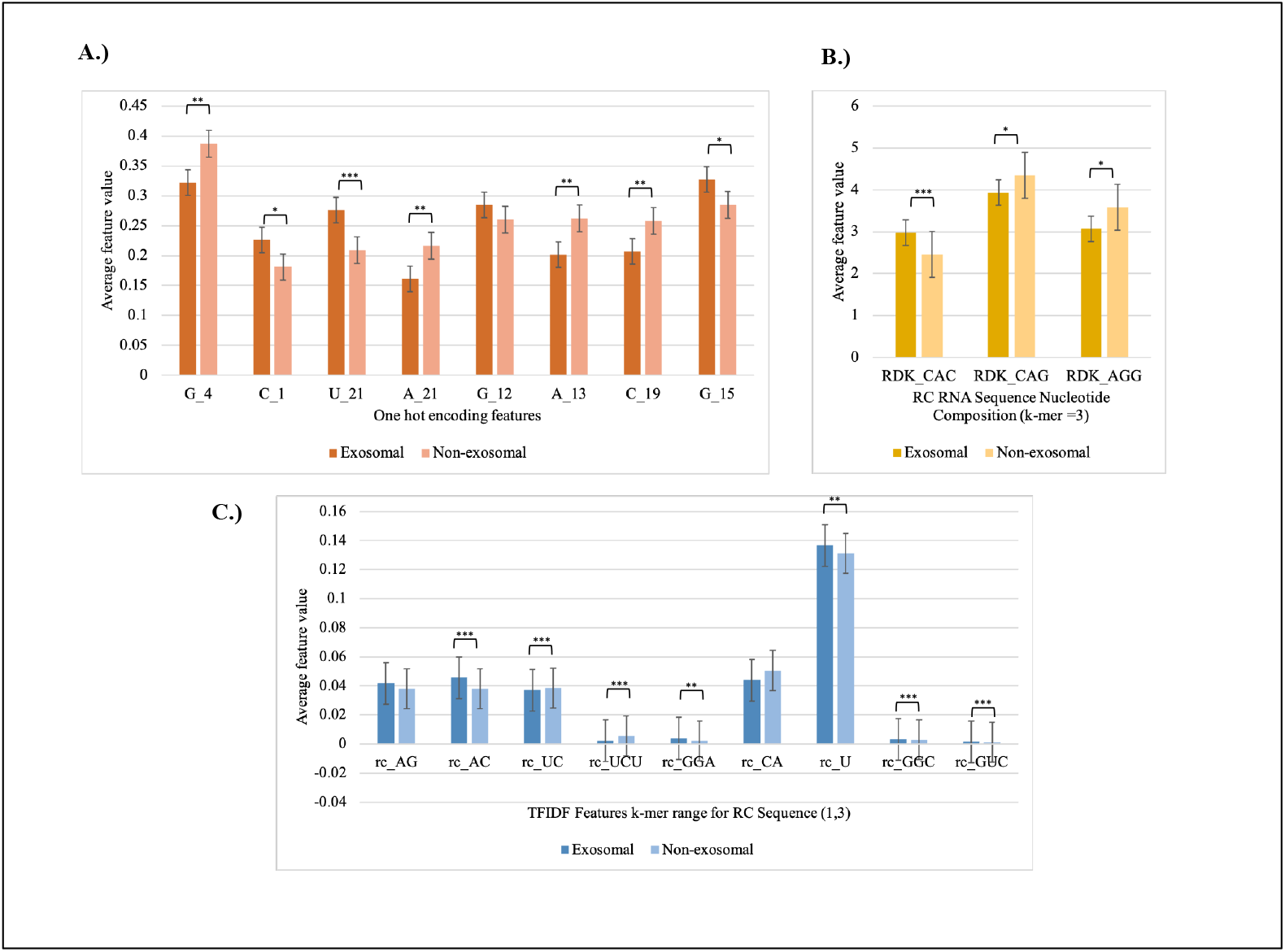
The 20 most important features for the classification of exosomal and non-exosomal miRNA

### 4. Comparison with existing methods

Presently, there are a few existing methods that predict the subcellular location of miRNA, with “exosome” being one of the locations. However, there is no tool that solely focuses on exosomal miRNA and predicts it with high accuracy. The tools that take miRNA sequences as input include miRNALoc, and EL-RMLocNet [50,51]. We wanted to compare the performances of these existing tools with our prediction tool. The tool miRNALoc reports the highest accuracy of 50% for predicting the subcellular localization of miRNA on an independent validation set, whereas EL-RMLocNet reports the highest AUC of 0.629 for predicting human miRNA on a benchmark dataset. To compare the results, we fed our independent validation set into the prediction servers, where we got an AUC of 0.494, an accuracy of 48.04%, and an MCC of -0.028 from the miRNALoc web server as compared to EmiRPred which gives an AUC of 0.73, accuracy of 67.62%, and MCC of 0.352. However, we were not able to get the predictions from EL-RMLocNet as it only takes one sequence at a time for the prediction, making it unfeasible to predict the validation set comprising 383 sequences.

### 5. Webserver and Standalone Software

One of the objectives of this study is to assist the scientific community working in the field of diagnostics or prognostics, specifically on exosomal miRNA-based biomarkers. This web server includes four modules: Predict, Design, Motif-scan, and BLAST-search. The “Predict” module accepts query sequences as input and predicts exosomal miRNA. Our “Design” module allows users to generate all possible mutant miRNA sequences and predict exosomal mutant miRNA. The “Motif-scan” module enables users to identify or scan known exosomal motifs in the query miRNA sequence. The “BLAST-search” module allows users to perform similarity searches against a database of known exosomal and non-exosomal miRNA sequences. This web server is compatible with smartphones, PCs, iMacs, and tablets. In addition to the web server, we have also created a Python package, a standalone tool, and a GitHub repository for this tool, available at the following links: https://webs.iiitd.edu.in/raghava/emirpred/ and https://github.com/raghavagps/emirpred.

## Discussions

Exosomes are small vesicles enclosed by a lipid membrane bilayer, secreted by most cells in the body. They carry various molecules, including proteins, lipids, metabolites, and RNA sequences [52]. In diagnostics, exosomes provide a non-invasive means to detect biomarkers for various conditions, such as cancer, neurodegenerative diseases, and cardiovascular disorders. In recent years a number of exosome-based biomarkers have been identified; for example, exosomal PD-L1 has been highlighted as a biomarker in non-small cell lung cancer [53]. Additionally, exosomal circRNAs have been implicated in colorectal cancer, offering potential as both diagnostic markers and therapeutic targets [54]. Exosomal miR-21 is frequently elevated in breast cancer, while exosomal miR-1246 has been linked to chemotherapy resistance in pancreatic cancer [55,56] In the realm of therapy, exosomes have been utilized to deliver CRISPR/Cas9 components for gene editing, showing efficacy in correcting genetic mutations in preclinical models of Duchenne muscular dystrophy [57]. Moreover, exosomes engineered to carry small interfering RNA (siRNA) targeting oncogenes like KRAS have demonstrated significant tumor suppression in pancreatic cancer models [58]. These advancements underscore the versatility of exosomes as both biomarkers and therapeutic delivery vehicles, paving the way for more personalized and effective medical interventions.

In this study, we focused on exosomal miRNA, the most abundant molecules found in exosomes, which hold immense potential as diagnostic and prognostic biomarkers. Our objective was to predict exosomal miRNA using various techniques, including motif identification, similarity search, machine learning, and deep learning. We identified several motifs and features that differentiate exosomal from non-exosomal miRNA. Initially, we discovered motifs or small patterns frequently found in exosomal miRNA. The most recurring motif was “GgapAAGCAC,” which appeared in 4 exosomal and 1 non-exosomal sequences in the validation set. Only a limited number of sequences contained these motifs, covering just 5.2% of miRNA sequences in the validation dataset. This indicates that motif identification alone is insufficient for predicting exosomal miRNA. Next, we employed a similarity search technique commonly used for sequence annotation. As mentioned in the results section, our similarity-based technique demonstrated a high probability of correct prediction but poor coverage. This suggests that alignment-based techniques (like motif and similarity search) are not adequate to predict all exosomal miRNA sequences.

In the last three decades, AI-based techniques have been heavily used to develop classification models in the field of computational biology, particularly in sequence classification. Thus, we utilized AI-based techniques to develop prediction models for this study. One of the advantages of AI-based models over alignment-based models is coverage of the dataset. Here, we systematically applied machine learning and deep learning techniques to develop prediction models. We also used all possible features, including compositional features, binary profiles, and embeddings, to develop models. In addition, we also developed large language models in this study to predict exosomal miRNA. Despite our best efforts, we achieved a maximum AUC of 0.707 with an MCC of 0.269 using AI-based models. Therefore, we developed an ensemble/hybrid method that combined the two techniques: Alignment-based models and AI-based models. Our ensemble method utilising these approaches achieved AUC of 0.73 with MCC 0.352.

It is important to compare newly developed methods with existing ones. To our knowledge, no method has been specifically developed for predicting exosomal miRNA sequences. However, we found miRNA subcellular localization methods like EL-RMLocNet and miRNALoc that can predict the subcellular localization of miRNA sequences. These subcellular localization methods predict multiple locations of miRNA in cells, including exosomal miRNA. Thus, we evaluated the performance of these methods on our independent validation dataset. As described in the results section, our method performed better than existing methods for exosomal miRNA. The limitation of this study is that, although the data is collected from experimentally validated studies, it is purely bioinformatics in nature. The features and motifs identified in this study have been previously reported to be associated with exosomal miRNA; however, in-depth studies with experimental validation are required.

## Author’s Biography

1. Akanksha Arora is currently pursuing a Ph.D. in Computational Biology at Department of Computational Biology, Indraprastha Institute of Information Technology, New Delhi, India.
2. Gajendra P. S. Raghava is currently working as a Professor and Head of Department of Computational Biology, Indraprastha Institute of Information Technology, New Delhi, India.

## Author’s contributions

AA collected and processed the data, implemented the algorithms, developed the prediction models, and built the front end and back end of the web server. AA, and GPSR prepared the manuscript. GPSR conceived and coordinated the project. All authors have read and approved the final manuscript.

## Acknowledgements

The authors are thankful to the Council of Scientific and Industrial Research (CSIR) for providing the fellowship. The authors are also thankful to the Department of Computational Biology, IIITD, New Delhi for its infrastructure and facilities. We thank the Department of Biotechnology (DBT) for providing an infrastructure grant to the institute (Grant BT/PR40158/BTIS/137/24/2021). We would like to acknowledge that figures were created using BioRender, and English was corrected using Grammarly.

## Data Availability Statement

All the datasets generated in this study are available at https://webs.iiitd.edu.in/raghava/emirpred/dataset.php and codes are available at https://github.com/raghavagps/emirpred.

